# Ontological Dimensions of Cognitive-Neural Mappings

**DOI:** 10.1101/524520

**Authors:** Taylor Bolt, Jason S. Nomi, Rachel Arens, Shruti G. Vij, Michael Riedel, Taylor Salo, Angela R. Laird, Simon B. Eickhoff, Lucina Q. Uddin

**Affiliations:** Department of Biomedical Engineering, Emory University, Atlanta, GA, USA; Department of Psychology, University of Miami, Coral Gables, FL, USA; Department of Neuroscience, Kenyon College, Gambier, OH, USA; Philips Research, Boston, MA, USA; Department of Physics, Florida International University, Miami, FL, USA; Department of Psychology, Florida International University, Miami, FL, USA; Institute of Systems Neuroscience, Medical Faculty, Heinrich Heine University Düsseldorf, Düsseldorf, Germany; Institute of Neuroscience and Medicine, Brain & Behaviour (INM-7), Research Centre Jülich, Jülich, Germany; Neuroscience Program, University of Miami Miller School of Medicine, Miami, FL, USA

**Keywords:** cognitive ontology, meta-analysis, fMRI

## Abstract

The growing literature reporting results of cognitive-neural mappings has increased calls for an adequate organizing ontology, or taxonomy, of these mappings. This enterprise is non-trivial, as relevant dimensions that might contribute to such an ontology are not yet agreed upon. We propose that any candidate dimensions should be evaluated on their ability to explain observed differences in functional neuroimaging activation patterns. In this study, we use a large sample of task-based functional magnetic resonance imaging (task-fMRI) results and a data-driven strategy to identify these dimensions. First, using a data-driven dimension reduction approach and multivariate distance matrix regression (MDMR), we quantify the variance among activation maps that is explained by existing ontological dimensions. We find that ‘task paradigm’ categories explain the most variance among task-activation maps than other dimensions, including latent cognitive categories. Surprisingly, ‘study ID’, or the study from which each activation map was reported, explained close to 50% of the variance in activation patterns. Using a clustering approach that allows for overlapping clusters, we derived data-driven latent activation states, associated with re-occurring configurations of the canonical fronto-parietal/salience, sensory-motor, and default mode network activation patterns. Importantly, with only four data-driven latent dimensions, one can explain greater variance among activation maps than all conventional ontological dimensions combined. These latent dimensions may inform a data-driven cognitive ontology, and suggest that current descriptions of cognitive processes and the tasks used to elicit them do not accurately reflect activation patterns commonly observed in the human brain.

## Introduction

The exponential growth of functional neuroimaging studies mapping constructs at the psychological level to patterns of brain activity (cognitive-neural mappings) has led to issues surrounding an adequate organizing ontology or taxonomy of these results ^1–4^. An ontology of a scientific domain serves both an organizational and descriptive function. In terms of organization, an ontology provides a standardized framework for managing, sharing and analyzing increasingly large databases of experimental analyses and data. Importantly, they also serve to describe experimental results in terms of natural divisions or dimensions of the phenomena under study. This function is particularly important for cognitive neuroscience, where a central goal is to map constructs at the psychological level to phenomena studied at a neurobiological level of analysis. The present goal of this study is to identify data-driven dimensions of a potential ontology based upon its ability to explain variability between patterns of brain activity.

Important progress in the development of an ontology for cognitive-neural mappings has been made using a combination of large-scale meta-analysis of task-based functional magnetic resonance imaging (task-fMRI) activation maps and topic modeling ^5–7^. At a general level, these studies map latent cognitive dimensions, derived from text documents ^5,6^ or task-paradigm descriptions ^7^, to patterns of brain activity. However, an important factor that needs to be addressed for an ontology of cognitive-neural mappings is what cognitive features of the task-fMRI environment drive the greatest differences in observed activation patterns. In other words, an adequate ontology of cognitive-neural mappings needs to identify those features of the task-fMRI experiment that explain the most variance among task-fMRI activation patterns.

To address this issue in a quantitative framework, we used a whole-brain representational similarity analysis (RSA) approach ^8^. Using this approach, we quantified the spatial similarity among whole-brain task-activation maps from the BrainMap database (Fox & Lancaster, 2002). Using multivariate distance matrix regression (MDMR; Zapala & Schork, 2006), we predict the variability between task-activation maps by existing ontological categories of the BrainMap database. The fundamental assumption of this approach is that the important features of the exiting ontology are identified in terms of the amount of variance they explain in task-activation map similarities.

Using the spatial similarity estimates among whole-brain task-activation maps, we also explore the possibility of estimating latent ontological dimensions directly from these similarities using a graph clustering approach. We take the opposite strategy of recent topic modeling approaches ^5–7^. In this approach, latent cognitive dimensions are estimated from experimental-descriptions and then projected onto patterns of brain activity ^5^, or co-occurring latent cognitive dimensions and patterns of brain activity are simultaneously estimated ^6,7^. In our approach, we reverse this chain of analysis. Commonly re-occurring whole-brain activation patterns, or activation states, are estimated directly from task-fMRI activation maps, and then projected onto experiment descriptions for interpretation. This approach assumes that the quality of a latent dimension is determined by how well these dimensions explain observable differences in task-fMRI activation patterns. Thus, latent cognitive dimensions are determined not in terms of their co-occurrence in text, but their co-occurrence with dominant whole-brain activation states. Importantly, these dimensions may or may not map cleanly onto existing latent cognitive dimensions.

## Results

### Outline of Approach

Our approach proceeds in three general steps (**Figure 1**). First, sparse non-negative matrix factorization (sparse-NMF) is applied to task-fMRI activation maps (N = 8919) reconstructed from experiment-reported peak-activation coordinates to derive a sparse ‘dictionary’ of brain networks. Spatial similarity estimates between activation maps could be computed at the voxel level, but this ignores spatial dependence between voxels. As opposed to other techniques (e.g., independent component analysis) for estimation of brain networks using the BrainMap database ^11,12^, sparse-NMF provides a data-driven network estimation technique uniquely suited for the data reported in the BrainMap database: all data are positive (i.e. positive activation values) and sparse (i.e. a small number of reported coordinates for each experiment). Sparse-NMF naturally leads to a parts-based parcellation of the brain given its enforcement of positive weights ^13^, even under low signal-to-noise conditions ^14^.

**Figure 1.**
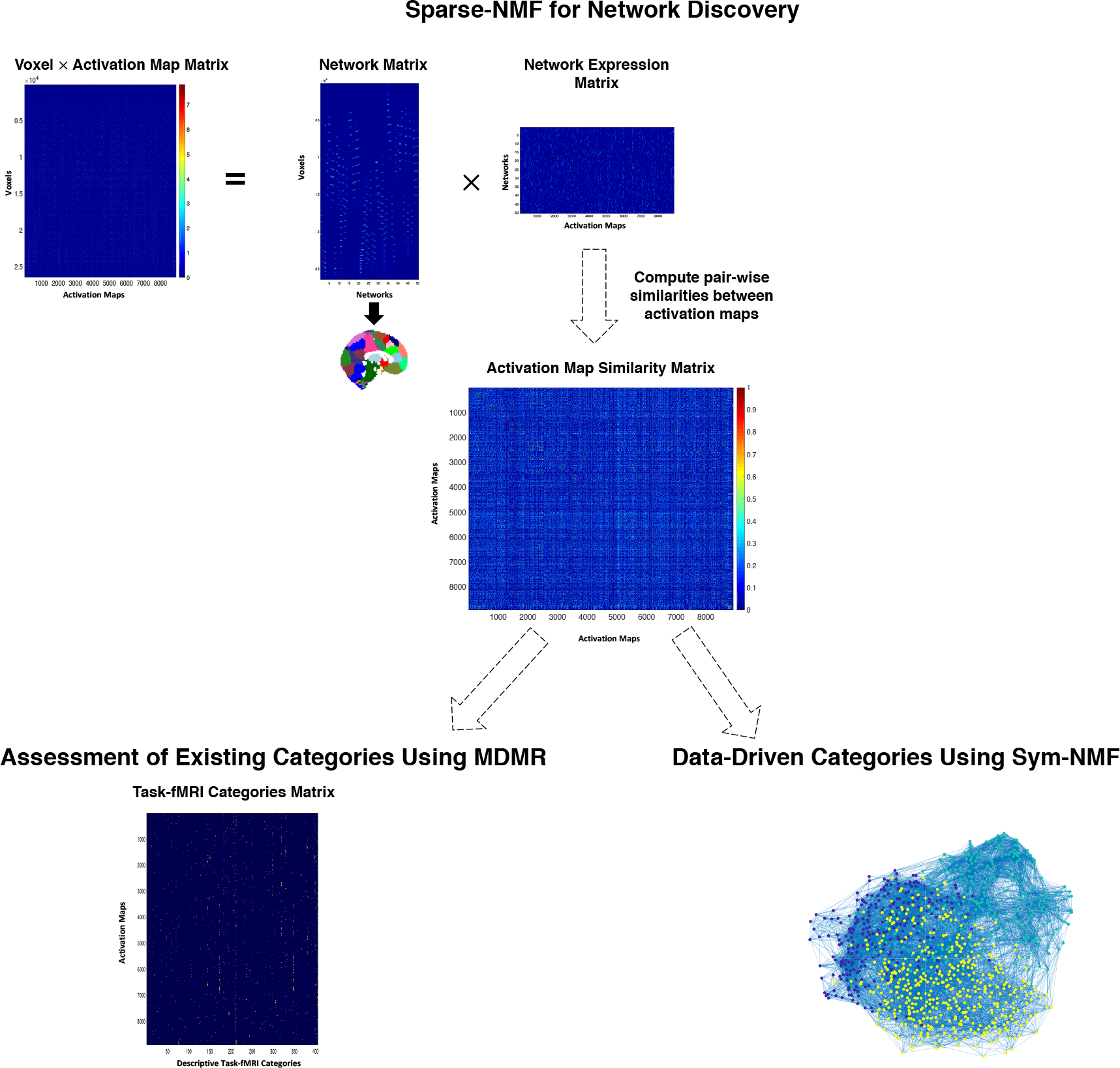
Illustration of Data-Driven Approach. First, latent network activation estimates are computed from activation maps derived from the BrainMap database using sparse non-negative matrix factorization (sparse-NMF). Sparse-NMF decomposes the activation map matrix into a ‘Network Matrix’ of network voxel weights, and a ‘Network Expression Matrix’ that reflects the activation estimates of all networks for each map. Next, an activation map dissimilarity matrix is constructed from dissimilarity estimates between latent network activation patterns for each map. With this computed dissimilarity matrix, we then assess existing ontological categories using MDMR, and derive potential data-driven ontological categories using Sym-NMF.

Second, we compute the spatial dissimilarity between the network activation estimates to a construct an activation map*activation map dissimilarity matrix. This can be seen as a large-scale whole-brain representational similarity analysis ^8^, where dissimilarity estimates are computed between large-scale activity patterns across the entire brain. We then predict the variability among activation maps from the experimenter-labeled ontological categories of the BrainMap database using multiple distance matrix regression (MDMR) ^10^. MDMR regresses a distance matrix onto a set of continuous or categorical predictors, and has extensive use in behavioral genomics ^15^ and studies of individual differences in resting-state functional connectivity (known as connectome-wide association studies) ^16^. The parameter of interest from this analysis is the explained variance estimate (R^2^) for five sets of ontological categories provided by the BrainMap database: analysis decisions, response type, stimulus type, task-paradigm class, and behavioral domain. These estimates provide information regarding what features of the existing ontology drive the greatest differences in task-activation maps.

Third, we convert the dissimilarity matrix into a similarity matrix for graph-based clustering. We use a graph clustering algorithm, known as symmetric non-negative matrix factorization (Sym-NMF), that has been shown to outperform commonly-used spectral graph-clustering algorithms ^17^. Importantly, Sym-NMF allows for overlapping communities of activation maps. As noted above, the goal of this clustering analysis was to estimate potentially overlapping whole-brain activation states. These states may prove useful as candidates for latent dimensions of a new data-driven ontology of cognitive-neural mappings. All code used in this manuscript are provided on the following webpage: https://github.com/tsb46/BrainMap-Cognitive-Ontology (**currently a work in progress**).

### NMF Results

In order to estimate the latent network activation estimates underlying each activation map, we applied a sparse-NMF algorithm to the full sample of 8919 task-fMRI activation maps. We derived a high-resolution network parcellation of 70 networks, the same high-model order estimated in previous analyses of the BrainMap database ^11,18^. The spatial similarity of activation maps replicated across alternative network solution sizes, as observed by the correlation between dissimilarity estimates of a 70-network solution size with smaller and larger solutions: N = 55, r = 0.91; N = 60, r = 0.93; N = 65, r = 0.96; N = 75, r = 0.95; N = 80, r = 0.94; N = 85, r = 0.92. Visual inspection of the 70-network solution (**Figure 2**) revealed a sparse parcellation of cortical and sub-cortical regions of the brain, corresponding to functionally relevant brain areas.

**Figure 2.**
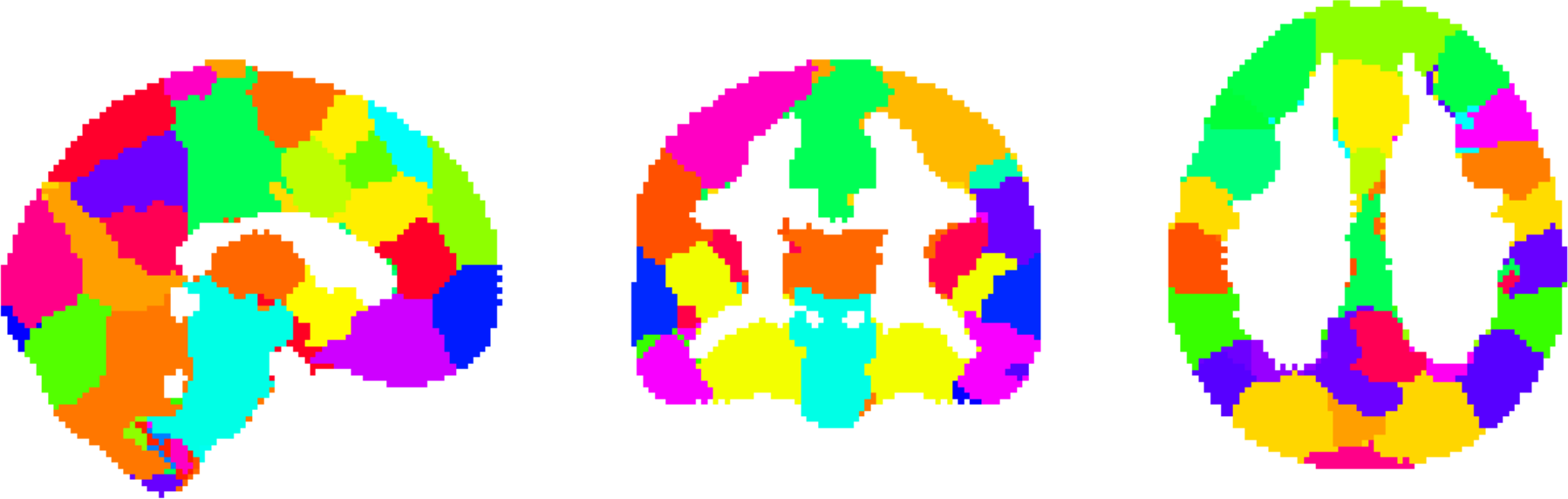
70-Network Parcellation. Visualization of the 70-network solution derived from the sparse-NMF algorithm. Networks are visualized with separate random colors.

### Prediction of Activation Map Dissimilarity from Existing Ontological Categories

Using the latent network activation estimates of the 70 networks we computed the activation map × activation map dissimilarity matrix with the cosine distance metric, representing the dissimilarity in latent network activation estimates between all activation maps. We used MDMR to regresses this dissimilarity matrix onto a set of ontological categories associated with each task-activation map. Categories provided by the BrainMap database fell into five domains: analysis decisions (subtraction vs. baseline contrast, and reported de-activation vs. activation), stimulus type (e.g. visual words, auditory stories, fixation cross, etc.), 3) response type (button press, speech, finger tapping, etc.), 4) paradigm class (encoding, task-switching, counting, etc.), and 5) behavioral domain (action execution, language semantics, working memory).

The BrainMap database contains nested data such that multiple task-activation maps are reported for a single study. To correct for this nesting in the BrainMap database, individual explained variance estimates for each categorical domain were corrected using a permutation test that respected study ID during permuting of task-activation map labels. Computation of the explained variance associated with each of the five categorical domains was carried out in a leave-one-out fashion. Each categorical domain was added to the remaining set of domains to compute the unique explained variance associated with each categorical domain.

The explained variance accounted for by all categorical domains (N_predictors_ = 603) in the model was 19.67%. The explained variance accounted for by analysis decisions (subtraction vs. baseline contrasts, and reported activation vs. de-activations), controlling for other domains, was 0.49% (*p* = 0.001). The explained variance accounted for by stimulus type, controlling for other domains, was 3.1% (*p* = 0.001). The explained variance accounted for by response type, controlling for other domains, was 0.57% (*p* = 0.014). The explained variance accounted for by paradigm class, controlling for other domains, was 4.17% (*p* = 0.001). The explained variance accounted for by behavioral domain, controlling for other domains, was 3.71% (*p* = 0.001). In summary, all categorical domains, which correspond to traditional ontological categories commonly used to describe cognitive-neural mappings, accounted for less than 20% of explained variance in activation maps.

We also ran a second model using only study ID as a predictor and we found that study ID, the study from which the task-activation map was reported, accounts for 44.71% of the variance in task-activation map differences. The single predictor of study ID accounted for more than twice the variance of the five categorical domains provided from the BrainMap database.

### Identification of Re-occurring Whole-Brain Activation States

As the full set of available ontological categories explain a limited amount of variance between whole-brain activation patterns, we next explore whether differences in whole-brain activation patterns can be explained by a smaller set of latent whole-brain activation states. We first apply a graph clustering approach, symmetric non-negative matrix factorization (Sym-NMF), to derive overlapping clusters of task-activation maps. Overlap allows for the possibility that a single task-activation map may represent a combination of multiple whole-brain activation states. We then project these latent dimensions onto the task-descriptive categories for behavioral and cognitive interpretation.

To choose the optimal number of clusters, we used a cross-validation approach to identify cluster sizes that exhibited stable cluster centroids across split-half samples. We repeated the two-fold cross-validation procedure for cluster sizes ranging from 2 to 20. Examination of average cluster centroid stability across cross-validation samples revealed two strong local minima of stability at a cluster sizes of four (*M_stability_*= 0.043) and seven (*M_stability_*= 0.055). Thus, further analyses were carried out on these cluster solutions.

In order to provide a functional interpretation of the clustering results, we computed the centroid activation maps and the prevalence of BrainMap ontological categories in each cluster. The centroid activation maps represent the weighted average activation pattern of the activation maps within the cluster. The association of a BrainMap category with a cluster was computed as the probability that BrainMap category appears given the cluster, normalized by the base probability of the cluster (see *Method and Materials*).

For the four-cluster solution, the centroid activation maps corresponded to functionally relevant brain areas (**Figure 4**). The clusters included a visual/dorsal-parietal activation pattern (Four Cluster 1; 4C1), fronto-parietal activation pattern (4C2), a default mode network (DMN) activation pattern (4C3), and an auditory/motor activation pattern (4C4). Consistent with previous reports of domain-general BOLD activation in anterior insula (AI) and dorsomedial prefrontal cortex (DMPFC) ^19^, activation in the AI and thalamus was observed across all centroid activation patterns. The BrainMap categories associated with the four clusters are consistent with previous functional descriptions of these brain areas. 4C1 was associated with paradigms requiring viewing and responding to or manipulating visual objects, including ‘mental rotation’, visual shape perception, and ‘spatial cognition’. 4C2 was associated with paradigms requiring a variety of complex high-level actions and behaviors, including ‘working-memory,’ ‘action inhibition’ and ‘language processing’. 4C3 was associated with paradigms requiring recall/recognition of previous information, inferences regarding others’ behavior (‘social cognition’), and various affective states (e.g. sadness, fear, reward, etc.). In addition, consistent with a DMN activation pattern, the activation maps within 4C3 were more likely to be reported as ‘de-activations’. 4C4 was associated with paradigms requiring overt/covert motor responses, along with listening or responding to auditory stimuli.

**Figure 3.**
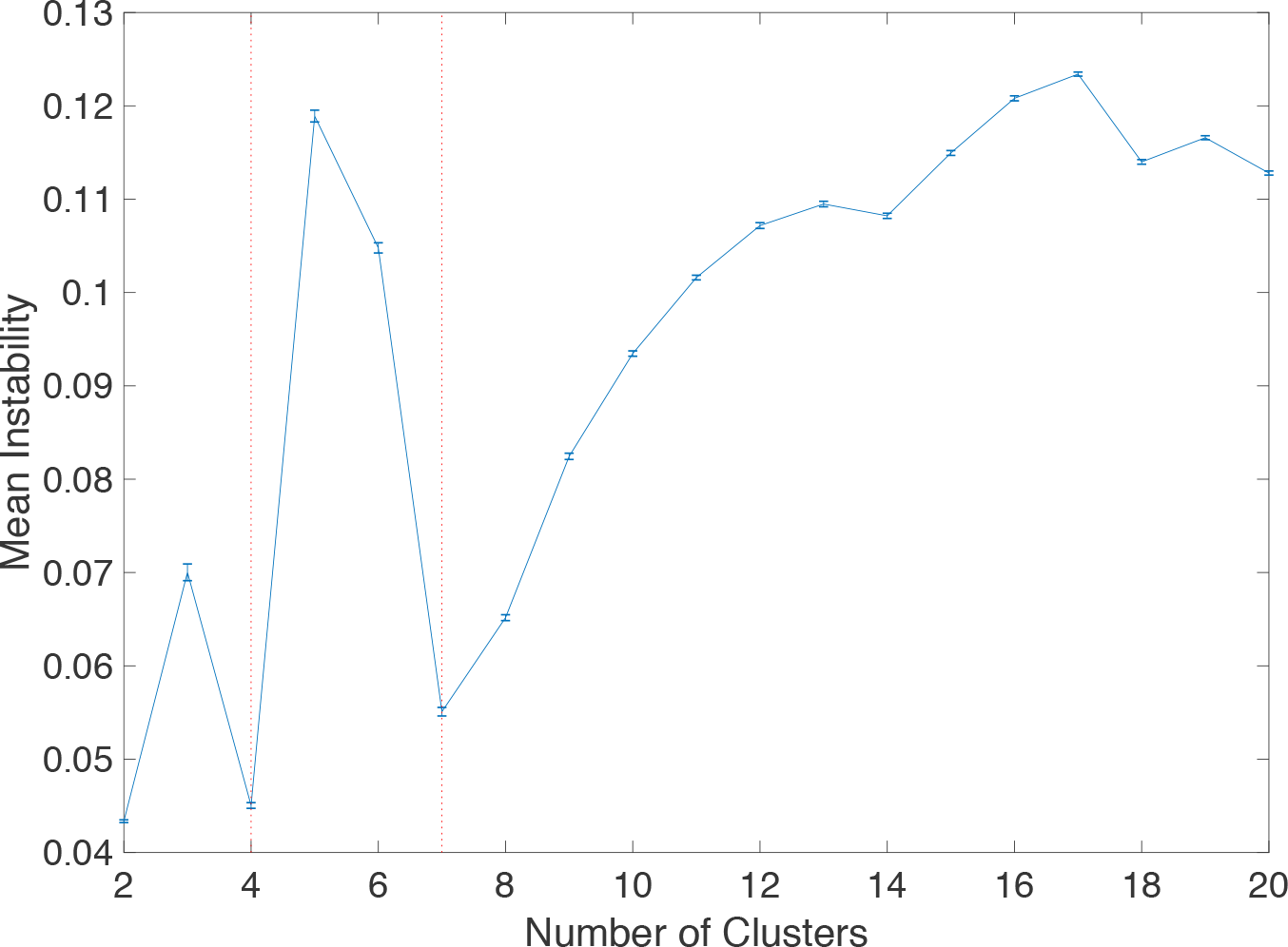
Instability Across Cluster Number. Mean instability (with standard error bars) across 100 random split-half samples for cluster numbers ranging from 2 to 20. Strong global minima were present at a cluster of four and seven (indicated by a vertical red line).

**Figure 4.**
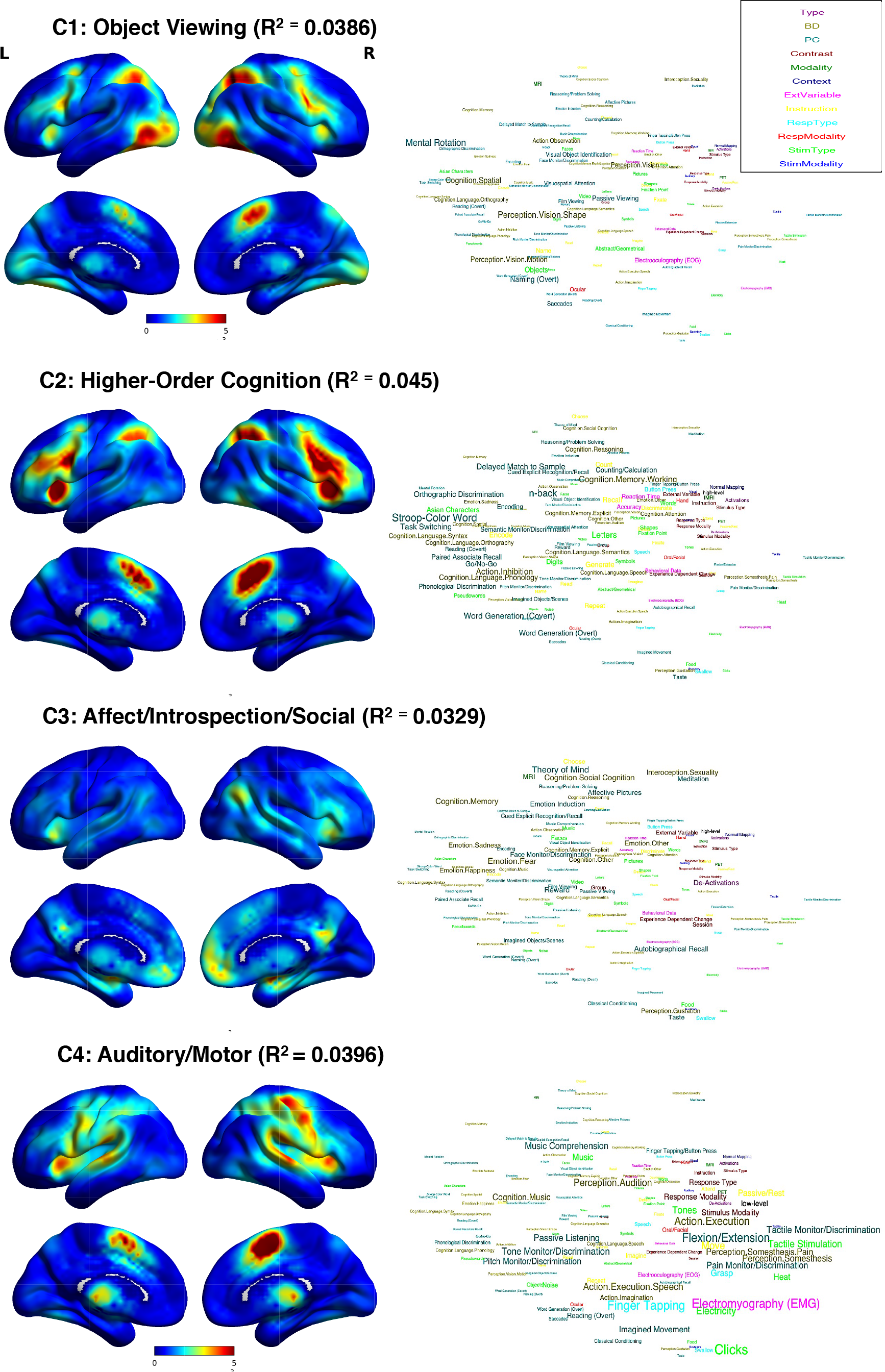
Centroid Activation Maps and Behavioral Decoding for Four-Cluster Solution. (BD = Behavioral Domain; PC = Paradigm Class; ExtVariable = External Variable; Stim = Stimulus; Resp = Response). Centroid activation maps with preliminary interpretative descriptions, and word cloud visualizations (presented to the right each centroid activation map) for the four-cluster solution. Experimental descriptors were sized by the degree of association with each cluster and their positions with respect to each other was determined by their ‘semantic closeness’. Experimental descriptors were colored according to the metadata category they belong to (e.g. Stimulus Modality, Behavioral Domain, etc.). A color key for each metadata category is provided in the top-right of the figure. The centroid activation maps represent the consistency of activation reported in each voxel across its cluster members (warmer colors represent a greater number of activations recorded at the voxel across activation maps for that cluster). Alongside each cluster label is the amount of unique explained variance that cluster explains in the similarity BOLD activation patterns, controlling for the remaining clusters.

The seven-cluster solution presented a slightly more fine-grained partition of the four-cluster solution. While 4C2 remained stable from the four-cluster to seven-cluster solution (7C7), the other three clusters from the four-cluster solution were split into two separate clusters in the seven-cluster solution. 4C4 in the four-cluster solution was split into two clusters with activation predominantly in sensory/motor cortices (7C2) and auditory cortices (7C5). Consistent with previous functional descriptions of these areas, the BrainMap categories associated with 7C2 included ‘action execution’, flexion/extension and tactile stimulation, and passive listening and pitching monitoring/discrimination for 7C2.

4C3 in the four-cluster solution was split in the seven-cluster solution into two clusters with activation predominantly in the default-mode network (7C4) and sub-cortical structures (basal ganglia and amygdala; 7C6). Consistent with previous functional descriptions of these areas, the categories associated with 7C4 included ‘theory of mind’, ‘social cognition’, and autobiographical recall, and affective stimuli and long-term memory for 7C6.

4C1 in the four-cluster solution was split in the seven-cluster solution into visual (7C1) and parietal-frontal clusters (7C3). Activation in 7C1 was predominantly observed in the visual cortex, with slightly weaker activation in the dorsal-parietal and frontal cortices. Activation in 7C3 was predominantly observed dorsal-parietal and frontal cortices, with slightly weaker activation in the visual cortex. Categories associated with 7C1 included passive viewing paradigms, including ‘action observation’, and visual shape perception, and perception of moving objects. Task-descriptive categories associated with 7C3 included active viewing paradigms, including ‘mental rotation’, ‘spatial cognition’, and delayed match to sample tasks.

An important observation apparent in the clustering solution is that BrainMap categories ‘distant’ in semantic space (farther apart in the word cloud) load strongly onto single activation states. In other words, categories that are semantically ‘distant’, are often not so ‘distant’ in the latent BOLD activation space. For example, consider the following two task paradigms: the Stroop color word task, and an overt word generation task. The psychological processes generally inferred to underlie these tasks include action inhibition ^20^ for the Stroop color word task, and lexical processing ^21^ for a word generation task. However, the whole-brain activation patterns of these paradigms are similar, as they both load strongly onto the same latent whole-brain activation state: 4C4 in the four-cluster solution and 7C7 in the seven-cluster solution. This suggests the neural response measured by task-fMRI is similar for the Stroop color word task and word generation task. Thus, whole-brain activation states may be informative in terms of mapping the similarity in functional anatomy between different cognitive or behavioral processes.

**Figure 5.**
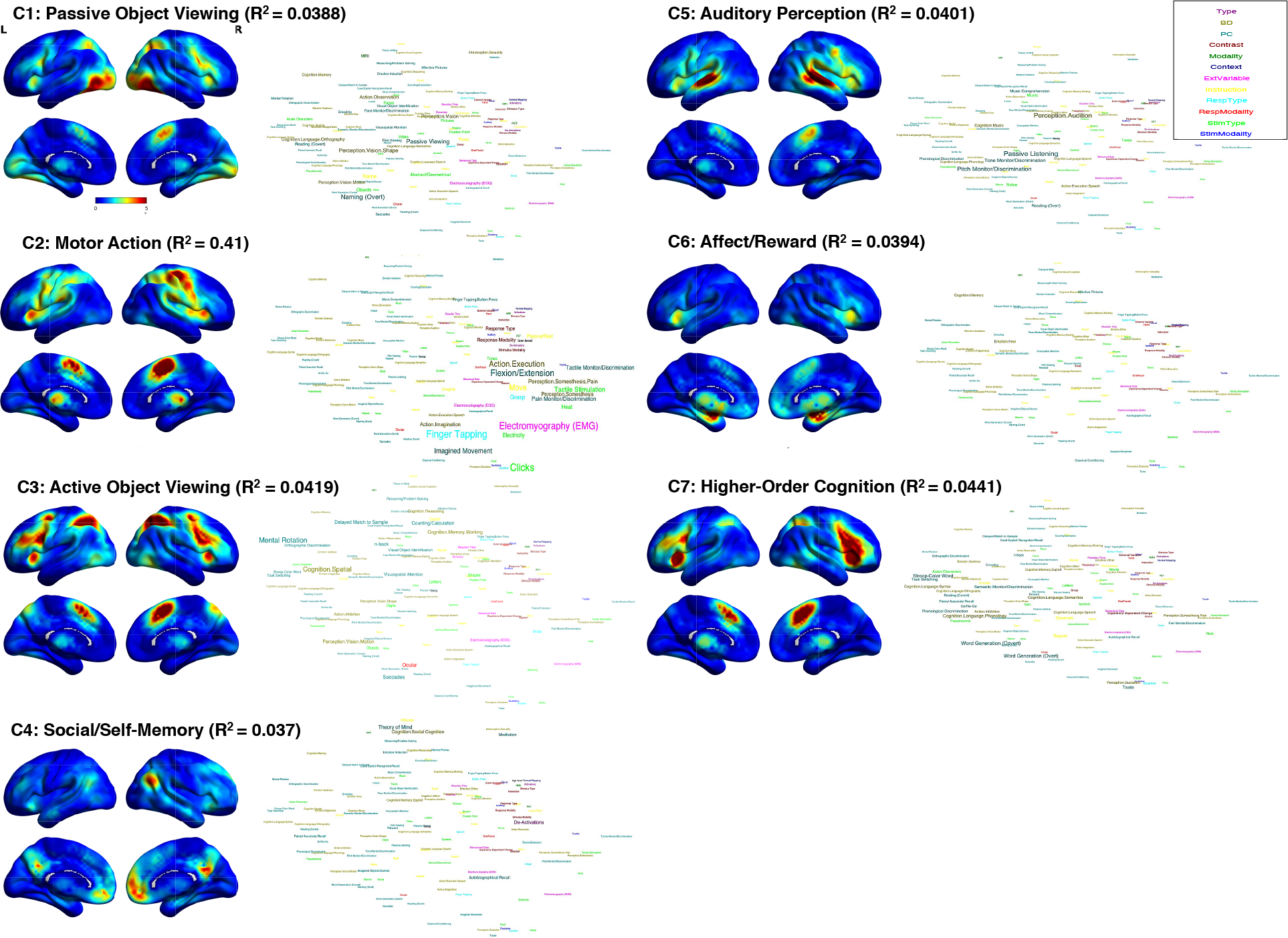
Centroid Activation Maps and Behavioral Decoding for Seven-Cluster Solution. (BD = Behavioral Domain; PC = Paradigm Class; ExtVariable = External Variable; Stim = Stimulus; Resp = Response). Centroid activation maps with preliminary interpretative descriptions, and word cloud visualizations (presented to the right each centroid activation map) for the four-cluster solution.

### Whole-Brain Activation State and BrainMap Category Comparison

To compare the explained variance from these latent whole-brain activation states to the existing ontological categories provided by the BrainMap database, we conducted an MDMR analysis regressing the weighted membership coefficients of the four- and seven-cluster separately onto the activation map * activation map dissimilarity matrix. The total explained variance for the four-cluster solution and seven-cluster solution were 20.95% (*p* = 0.001) and 34.37% (*p* = 0.001), respectively. Thus, both lower-order latent activation states (N =4 and N = 7) explained a greater amount of variance in task-activation map dissimilarity than the full model of existing ontological categories (N = 603). This difference in explained variance can be visualized with a re-ordered activation map*activation map similarity matrix according to either the latent activation states or two example categorical domains (*Behavioral Domain* and *Paradigm Class*; **Figure 5**).

**Figure 5.**
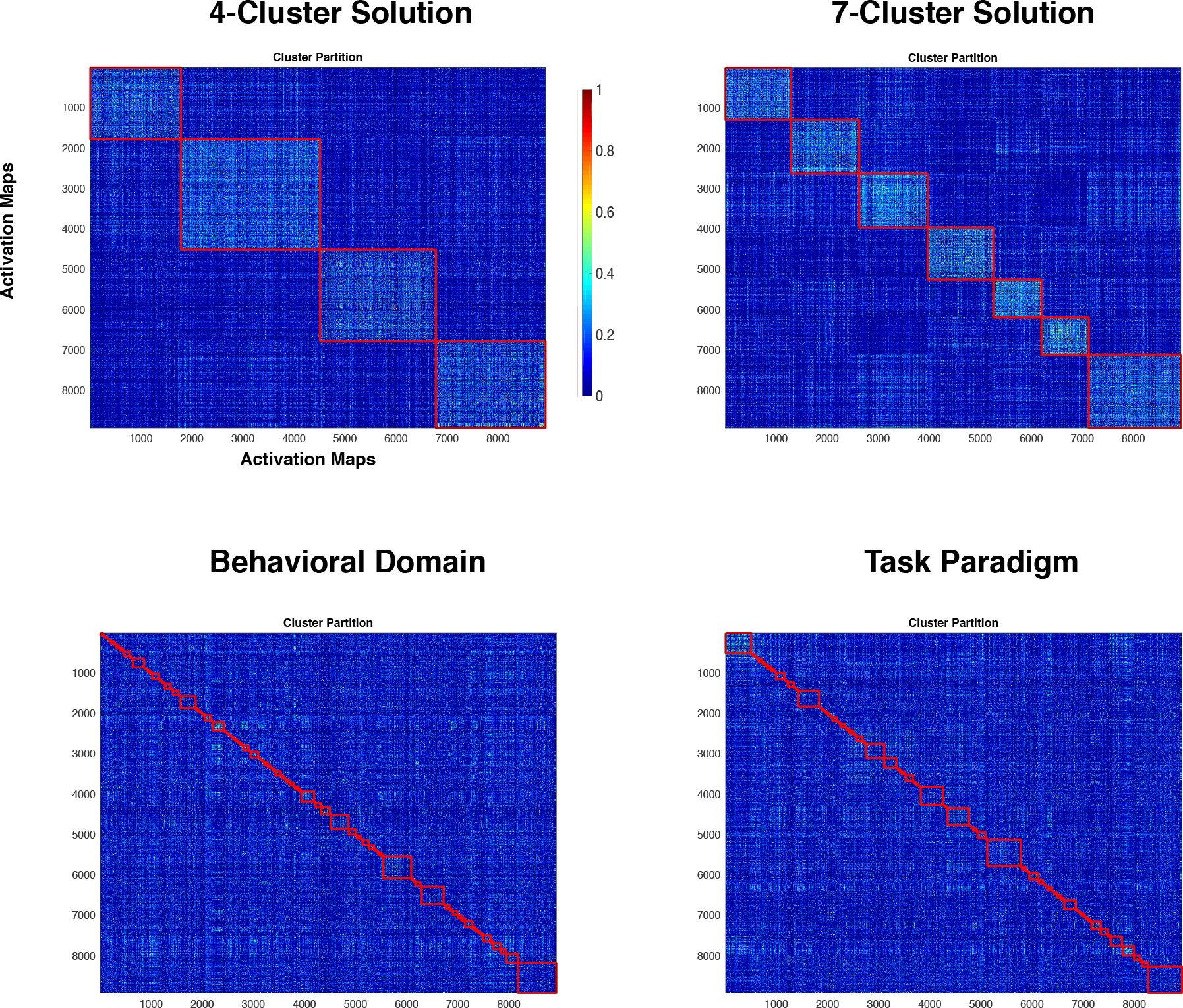
Comparison of Latent Activation State Clusters and Task-Descriptive Category Clusters. To visualize the differences in explained variance between the latent activation states and two task-descriptive categories (*Behavioral Domain* and *Paradigm Class*), we visualized a re-ordered activation map * activation map similarity matrix according to the latent activation state and task-descriptive categories. The re-ordered similarity matrices for the latent activation state and task-descriptive categories were created by re-organizing each activation map in the similarity matrix according to their dominant whole-brain activation state cluster (i.e. assignment of each map to the cluster with the strongest loading) or *Behavioral Domain/Paradigm Class*. The values in the similarity matrices vary from 0 (no similarity; cool colors) to 1 (max similarity; warm colors). The red lines organized along the diagonal of the similarity matrix trace groups of activation maps that belong to a single cluster according to their latent activation state or *Behavioral Domain/Paradigm Class* category. Similarity matrices with brighter colors within the red cluster lines, and cooler colors outside, correspond to more compact and well-separated groups of task-activation maps.

## Discussion

A driving goal of cognitive neuroscience is to generate and constrain theories and models of cognitive processes using functional neuroimaging data. Directed at this goal, the primary aim of this study was to identify the dominant cognitive dimensions upon which whole-brain blood-oxygen-level dependent (BOLD) activation maps vary. In other words, what dimensions of the task-fMRI experiment is the BOLD response most attuned do? Task-based fMRI research has conventionally used a fluid set of ontological categories to interpret the differences between activation maps. Using a novel analysis pipeline applied to the BrainMap database, we quantify the variance explained by these existing ontological categories, describing various features of the task-fMRI experiment. The full model of existing ontological categories (N = 603 predictors) explained a moderate proportion of variance (R^2^= 0.197) in the similarities between activation maps. We find that ‘stimulus type’, ‘paradigm class’ and ‘behavioral domain’ uniquely explain a small (R^2^< 0.05), but non-trivial amount of variance in similarity between activation maps.

It is difficult to assess these explained variance estimates in terms of absolute value, as the disparate quality of the activation maps (derived from peak-activation coordinates) place an unknown upper limit on the variance explainable by the task-descriptive categories. However, we can make relative comparisons. We find that the observable domain of ‘paradigm class’ (or task paradigm), explains the greatest amount of variance among the task-descriptive categories, controlling for other domains, (R^2^ = 0.0417). This suggests that differences in task experiments drive the most observed differences in fMRI task-activation patterns over and above differences in other domains (e.g. behavioral domain). Thus, researchers should be aware that the choice of experimental task may be of more crucial importance than whether they are thought to be eliciting the same purported cognitive or behavioral process. Surprisingly, a substantial portion of variance in the similarity between activation maps can be explained by the study ID (R^2^ = 0.447); over two times greater than the full set full model of task-descriptive categories (R^2^ = 0.197). This can be for multiple reasons: 1) differences in peak-coordinate reporting, 2) differences in pre-processing pipelines and scanner hardware, or 3) differences in significance thresholding procedures ^22^. These differences in study ID may be reduced with the sharing of unthreshold statistical maps ^23^, that avoids peak-coordinate reporting and thresholding procedures ^19^.

We assume that the quality of latent cognitive or behavioral descriptors for a task-fMRI ontology is contingent upon how well these descriptors explain differences in BOLD activation patterns. Thus, we attempt to derive a set of latent activation states and associated cognitive and behavioral descriptors that maximize the explained variance between BOLD activation patterns. With just a set of four (or seven) latent activation states with sensible cognitive interpretations, we can explain a greater amount of variance in BOLD activation maps than the full set of ontological categories of the BrainMap database (N = 603). One might argue that this is trivial: one would expect that a solution built to maximize explained variance between activation patterns will of course explain more variance than the *a priori* ontological categories. However, we do not believe this is trivial for two reasons: 1) though it is trivial fact that all clustering algorithms, at a general level, estimate clusters that maximize explained variance, it is not trivial that a *low-dimensional* clustering solution (N= 4 or N = 7) can explain greater variance than the high-dimensional task-descriptive categories (N = 603). Relatedly, 2) explained variance of activation map dissimilarities *should* be our criterion for what constitutes an adequate ontology of cognitive-neural mappings. For example, if a four-cluster solution explained 99% of the variance in task-activation map dissimilarity, adding more ontological categories amounts to an unnecessary overparameterization of this space.

The BrainMap categories associated with the latent activation states from the 4-cluster and 7-cluster map onto plausible neurocognitive systems. The four latent activation states can be generally described as object-viewing (4C1), higher-order cognition (inhibition/control/language) (4C2), self-memory/affective/social (4C3), and auditory-stimulus/motor-action (4C4) states. The seven latent activation states can be generally described as passive object-viewing (7C1), sensory-motor (7C2), active object-viewing (7C3), social/theory of mind (7C4), auditory (7C5), affective (7C6) and higher-order cognition (7C7). Examination of the four-cluster and seven-cluster solutions centroid activation patterns revealed a set of regions commonly activated across the BrainMap database. Consistent activation was observed across regions that make up the traditionally described ‘task-positive’ and ‘task-negative’ activation/de-activation pattern ^19,24,25^. The ‘task-positive’ activation pattern traditionally includes elements of the fronto-parietal (DLPFC, SPC and DMPFC) and salience networks (dorsal anterior cingulate cortex and AI). Consistency of activation was particularly predominant in the AI across all centroid activation maps, in accord with previous studies demonstrating the ubiquity of AI activations across task paradigms ^26^. The ‘task-negative’ de-activation pattern, generally restricted to the DMN ^25^, was represented in both the four- and seven-cluster solution. Consistent with the ‘task-negative’ ascription, the activation maps of these clusters were more likely to be reported as ‘de-activations’.

### Limitations

One objection to the current approach is that it is circular in some sense. According to this objection, the use of conventional task-descriptive categories of cognitive functioning and task paradigms for interpretation in the behavioral decoding analysis essentially reifies these categories in our neural-driven categorization. Thus, the analysis relies on a circular inference. In response, we agree that the mix of ordinary and technical terms codified in the BrainMap experimental descriptors is relied upon in our descriptions of the data-driven categories, but reliance on this terminology does not constitute circularity. The behavioral decoding analysis maps clusters to their most representative experimental descriptors, but does not *identify* the clusters with those experimental descriptors. For example, descriptions of complex physics topics often rely on ordinary linguistic terminology, but this description does not then identify the physical systems themselves with these ordinary terms. In the same manner, describing the data-derived categories in terms of their association with ordinary or current technical terms of cognitive science does not constitute circularity. By examining the pattern of experimental descriptors associated with each cluster, we may find pointers to an underlying neural process that unifies these descriptors.

Due to the disparate representation of activation maps in the BrainMap database in terms of peak-activation coordinates, the precision of the distinctions between possible clusters is limited. However, the substantial sample size of reported task-fMRI experiments provided by the BrainMap database is unmatched in terms of power and detailed experimental metadata. The increasing size of databases containing unthresholded activation maps, such as NeuroVault ^23^, offers the potential for a more precise or fine-grained categorization using the approach applied in this study. In addition, the task-activation maps in the BrainMap database were computed using a variety of analytic approaches and experimental designs, which introduces an extra source of variability among the activation maps that cannot be accounted for in our analysis. Nevertheless, we hope the results presented here provide a starting point for a well-developed ontology of cognitive-neural mappings.

## Acknowledgements

Peter Fox/Brainmap. This work was supported by award R01MH107549 from the National Institute of Mental Health to LQU and award 1631325 from the National Science Foundation to ARL.

## Methods and Materials

### Construction of Activation Maps from BrainMap Coordinates

At the time of the analysis, the BrainMap database ^9^ contained 15900 experimental contrasts from 3216 published manuscripts. Associated with each experimental contrast were the coordinates of statistically significant peak-activation coordinates in MNI152 and Talairach coordinate spaces. Coordinates in Talairach space were converted to MNI152 coordinates using the transform described in Lancaster et al. ^27^. The number of contrasts was reduced to ensure that only experimental contrasts of interest were included in the analysis. The only type of experimental contrast included in the analysis was a ‘normal mapping’ experiment, or exclusively a contrast within healthy participant groups (e.g. no drug treatment). These criteria resulted in a total number of 8919 experimental contrasts for the current analysis.

For each of the 8919 experimental contrasts, modeled activation maps were constructed by placing a 12-mm FWHM Gaussian kernel around the center of each peak-activation coordinate reported for each contrast. This kernel size is identical to a previous analysis of the BrainMap database ^11^. Because the modeled activation maps are extremely sparse (i.e. consisting mainly of zeros), they were sub-sampled by a power of 2 along the X, Y and Z directions for computational feasibility. This sub-sampling resulted in 26459 voxel values per activation map.

### Network Discovery using Non-Negative Matrix Factorization

To estimate the latent network activation estimates associated with each activation map, we utilized sparse-NMF. The activation maps were first vectorized and placed into a 26459 (voxel) 8919 (activation map) matrix. This was then input to the sparse-NMF algorithm to model the voxel × activation map matrix as the multiplicative combination of a network matrix and network expression matrix. Each element of both matrices is constrained to be >= 0. The network matrix (voxel × selected number of networks) contains the voxel weights of each network. The network expression matrix (selected number of networks × activation map) represents the latent network activation estimate of each network for each activation map. The NMF algorithm implements the following minimization objective:

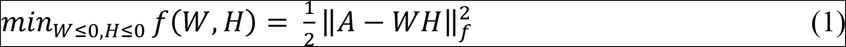

where A is the voxel × activation map matrix, W is the network matrix, H is the network expression matrix, and ‖ ‖*_f_* is the Frobenius norm of the residual of the W*H matrix projected onto the A matrix.

The conventional estimation algorithm for NMF estimates elements of the network and network expression matrix by minimizing the above objective using a multiplicative update rule ^13^, which is computationally infeasible given the dimensionality of the input matrix (26459 × 8919 matrix). Thus, we used a fast NMF estimation procedure using the block principal pivoting/active set method described by Kim and Park ^28^ and implemented in MATLAB code provided on their webpage (https://www.cc.gatech.edu/~hpark/nmfsoftware.php). We used a variation of NMF, sparse-NMF, that adds an additional sparsity constraint imposed on the network matrix (W matrix) by adding the L1 Norm of the sum of each column of the network matrix to the minimization objective above (Eq. 1). The estimation algorithm searches for a *local* minimum, and is thus initialization-dependent. Rather than using a random initialization of the network and network expression matrix, we use a singular-value decomposition (SVD) of the voxel × activation map matrix as the starting initialization of the algorithm described by Boutsidis and Gallopoulos (2008), and implemented by Sotiras et al. ^30^. The code implemented was provided on the following webpage: https://github.com/trigeorgis/Deep-Semi-NMF/blob/master/matlab/NNDSVD.m.

The network expression estimates were used in the subsequent computation of task-activation map dissimilarity. We chose a higher-order network solution of 70 networks, comparable to previous analyses of the BrainMap database ^11,18^. However, the spatial dissimilarity estimates replicated across alternative network solution sizes, as observed by the correlation between dissimilarity estimates of a 70-network solution size with smaller and larger solutions: N = 55, r = 0.91; N = 60, r = 0.93; N = 65, r = 0.96; N = 75, r = 0.95; N = 80, r = 0.94; N = 85, r = 0.92.

### Prediction of Activation Map Dissimilarity with Task-Descriptive Categories

#### Dissimilarity Matrix Computation

Using the latent network activation estimates of the 70 networks we computed the activation map × activation map dissimilarity matrix with the *cosine* distance metric, representing the dissimilarity in latent network activation estimates between all activation maps. The cosine distance metric was used over other possible distance metrics because it is advantageous for high-dimensional and sparse data ^31,32^. The pair-wise dissimilarity between two activation maps in network expression profiles could vary from 0 (perfect similarity) to 1 (no similarity).

#### BrainMap Categories Predictor Matrix Preprocessing

Ontological categories of the BrainMap database fell into five domains: 1) analysis decisions (subtraction vs. baseline contrast, and reported de-activation vs. activation), 2) stimulus type (e.g. visual words, auditory stories, fixation cross, etc.), 3) response type (button press, speech, finger tapping, etc.), 4) paradigm class (encoding, task-switching, counting, etc.), and 5) behavioral domain (action execution, language semantics, working memory). For details of each category, please see: http://www.brainmap.org/taxonomy/. For each domain, categories were dummy-coded leaving the following variables as the reference variable for each domain: ‘words’ (stimulus type), ‘write’ (response type), ‘n-back’ (paradigm class), and ‘Perception.Vision.Shape’ (behavioral domain).

Because the BrainMap categories (excluding analysis decisions) allow multi-label classification (i.e. activation maps can have more than one sub-category), multi-label classifications were treated as separate clusters. For example, an activation map classified as Action Inhibition and Attention was treated as belonging to an Action Inhibition/Attention cluster, as opposed to the belonging to both an Action Inhibition and Attention cluster. This increased the number of clusters, and allowed for more precise classifications. A small percentage of activation maps belonged to multi-label categories with fewer than five members (N = 435 for stimulus type; N = 61 for response type; N = 706 for paradigm class; N = 599 for behavioral domain), and these multi-label categories were re-categorized as ‘unclassified.’ Inclusion of multi-label classifications without a membership size cutoff resulted in a heavily overparameterized model (1508 variables), and large sets of perfectly multicollinear predictors (resulting in a rank deficient matrix).

Additional categories were also excluded from the analysis, including those categories with high collinearity with another category (*r >0.99*; 14 categories), ‘Not Defined’ categories, and multi-label categories that included a ‘None’ classification and a non-None classification. The final dummy coded matrix contained a total of 603 dummy coded variables, including 163 stimulus type variables, 29 response type variables, 215 paradigm class variables, 194 behavioral variables, and 2 analysis decision variables.

#### Multivariate Distance Matrix Regression

We used multivariate distance matrix regression (MDMR) ^10^ to model the variability in whole-brain activation map dissimilarity explained by the dummy-coded category predictor matrix. Computation of the unique explained variance associated with the five categorical domains was carried out in a leave-one-out fashion. Each domain of categorical predictors was added to the remaining set of categorical predictors to compute the unique explained variance associated with each categorical domain.

Activation maps reported in the BrainMap database have a nested structure: multiple activation maps are reported for the same study. To correct for this nesting in the BrainMap database, explained variance estimates for each categorical domain were corrected using a permutation test that respected study ID. The permutation test involved a nested reshuffling of activation map labels within each study set. After each reshuffling, an R^2^ estimate for each set of predictors was computed to construct a null distribution of R^2^ values. The p-value of the original R^2^ was then computed as the percentile of the original R^2^ in the null distribution. The permutation test was run 1000 times.

### Graph Clustering Approach

Symmetric non-negative matrix factorization (SymNMF) applies the non-negative matrix factorization framework to pair-wise similarity matrices ^17^. In the sparse-NMF algorithm described above, the voxel*activation map matrix is factorized into a multiplicative combination of a network matrix (W) and network expression matrix (H). In the symmetric-NMF framework, the symmetric activation map*activation map similarity matrix (S) is factorized into a cluster membership matrix (H) multiplied by itself. More formally, the symmetric-NMF minimizes the following objective function:

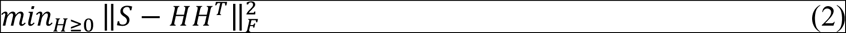

where S is the activation map × activation map similarity matrix, H is the cluster membership matrix, and ‖ ‖*_f_* is the Frobenius norm of the residual of the H*H^T^ matrix projected onto the S matrix. Importantly, SymNMF allows for each activation map to have non-negative weights on more than one cluster. Thus, an activation map can have varying degrees of membership for each cluster. The MATLAB code implement for SymNMF was provided on the following webpage: (https://github.com/andybaoxv/symnmf).

### Determination of the Number of Clusters

To choose the optimal number of clusters, we used a repeated two-fold cross-validation procedure for cluster sizes ranging from 2 to 20. First, 100 random split-half samples (n = 4459) of the total sample of activation maps were generated. Next, the above SymNMF algorithm was applied to all 100 split-half pairs. Finally, the average pair-wise instability between the 100 split-half cluster centroids was measured using an Amari-type quantity procedure developed by Wu et al. ^33^ (https://github.com/bdgp/staNMF). The average instability, which can vary from 0 (perfect stability) to 1 (no stability), was then plotted across all solutions to search for network solution sizes with low instability. Plotting of the average instability values across all solutions sizes from 2 to 20 revealed two strong local minima in average instability (**Figure 3**).

### Centroid Activation Patterns and Behavioral Decoding of Clusters

The resulting community partitions (i.e., clusters) at the two levels of resolution (c = 4 and c = 7) were interpreted in terms of their centroid activation pattern and the BrainMap task-descriptive categories. The centroid activation map of each cluster was computed in two steps: 1) network expression profiles of each activation map within the cluster were averaged to yield a centroid network expression profile, and 2) the centroid network expression profile was then projected onto voxel space to create the centroid activation map using the following equation:

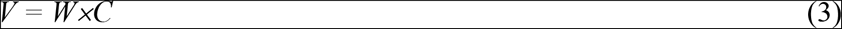

where *V* is 26459 (voxel) × 1 centroid activation map, *W* is the network matrix (from the NMF solution), and *C* is the centroid network expression profile. For visualization, the missing voxel values in the centroid activation map from the sub-sampling procedure described above were interpolated using a penalized least squares procedure ^34^ (https://www.mathworks.com/matlabcentral/fileexchange/27994-inpaint-over-missing-data-in-1-d--2-d--3-d--n-d-arrays). The resulting images were then smoothed using a 6mm FWHM Gaussian kernel.

Behavioral decoding of the clusters was performed using the task-descriptive categories used in the MDMR analysis above. In addition to Stimulus Type, Response Type, Paradigm Class, Behavioral Domain and Analysis Decisions, we used the following additional task-descriptive categories: Stimulus Modality, Response Modality, Instruction, and External Variable. Of note, some task-descriptive categories were labeled ‘None’ or ‘Unknown,’ and were excluded from the analysis. To behaviorally decode each cluster ***i*** for each task-descriptive category ***s*** of all metadata categories, we computed the forward inference probability normalized by the probability of that cluster:

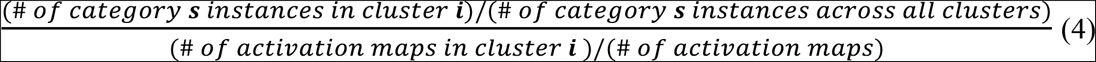

where the numerator represents the *Pr*(*Cluster | Category*) and the denominator represents the *Pr*(*Cluster*).

### Visualization of Behavioral Decoding

Given the number of task-descriptive categories assessed (*n* = 144; from the above-mentioned categories), we used a data-driven word cloud visualization to illustrate the results. The word cloud visualization displays text with varying sizes and colors throughout space in a visualization window. For each cluster, word locations were positioned by the ‘semantic closeness’ of each task-descriptive category. The size of the text was proportional to the behavioral decoding result from equation (4). To estimate the ‘semantic closeness’ of the sub-categories we did the following: 1) counted the appearances of all the sub-categories for each activation map to create a 144 (sub-category) × 8919 (activation map) sub-category count matrix, 2) estimated the 144×144 sub-category distance matrix by computing the pair-wise *Jaccard* distance in the counts between all pairs of sub-categories, and 3) projected the distances between all 144 sub-categories onto a two-dimensional subspace using a nonmetric multidimensional scaling (MDS) algorithm using the *mdscale* scale function in MATLAB (https://www.mathworks.com/help/stats/mdscale.html). Metadata categories were distinguished by different colors. In many cases, the original word cloud visualization output spatially overlapping sub-category terms. Thus, for clarity, we manually moved spatially overlapping sub-category terms to reduce overlap, while respecting their original position along the dimensions derived from the MDS solution. We used publically available code for displaying the word cloud visualization, provided on the following webpage: (https://www.mathworks.com/matlabcentral/fileexchange/53016-wordcloud--classical-).

The data that support the findings of this study are available from www.BrainMap.org, but restrictions apply to the availability of these data, which were used under license for the current study, and so are not publicly available. Data are however available from the authors upon reasonable request and with permission of BrainMap.

## References

1. Hastings, J. et al. Interdisciplinary perspectives on the development, integration, and application of cognitive ontologies. Front. Neuroinformatics 8, (2014).

2. Laird, A. R. et al. Neural architecture underlying classification of face perception paradigms. NeuroImage 119, 70–80 (2015).

3. Poldrack, R. A. et al. The cognitive atlas: toward a knowledge foundation for cognitive neuroscience. Front. Neuroinformatics 5, 17 (2011).

4. Turner, J. A. & Laird, A. R. The cognitive paradigm ontology: design and application. Neuroinformatics 10, 57–66 (2012).

5. Poldrack, R. A. et al. Discovering Relations Between Mind, Brain, and Mental Disorders Using Topic Mapping. PLoS Comput. Biol. 8, (2012).

6. Rubin, T. N. et al. Decoding brain activity using a large-scale probabilistic functional-anatomical atlas of human cognition. PLOS Comput. Biol. 13, e1005649 (2017).

7. Yeo, B. T. T. et al. Functional Specialization and Flexibility in Human Association Cortex. Cereb. Cortex N. Y. N 1991 25, 3654–3672 (2015).

8. Haxby, J. V. et al. Distributed and Overlapping Representations of Faces and Objects in Ventral Temporal Cortex. Science 293, 2425–2430 (2001).

9. Fox, P. T. & Lancaster, J. L. Mapping context and content: the BrainMap model. Nat. Rev. Neurosci. 3, 319–321 (2002).

10. Zapala, M. A. & Schork, N. J. Multivariate regression analysis of distance matrices for testing associations between gene expression patterns and related variables. Proc. Natl. Acad. Sci. 103, 19430–19435 (2006).

11. Smith, S. M. et al. Correspondence of the brain’s functional architecture during activation and rest. Proc. Natl. Acad. Sci. U. S. A. 106, 13040–13045 (2009).

12. Toro, R., Fox, P. T. & Paus, T. Functional Coactivation Map of the Human Brain. Cereb. Cortex N. Y. NY 18, 2553–2559 (2008).

13. Lee, D. D. & Seung, H. S. Learning the parts of objects by non-negative matrix factorization. Nature 401, 788–791 (1999).

14. Lohmann, G., Volz, K. G. & Ullsperger, M. Using non-negative matrix factorization for single-trial analysis of fMRI data. NeuroImage 37, 1148–1160 (2007).

15. Schmitz, R. J. et al. Transgenerational Epigenetic Instability Is a Source of Novel Methylation Variants. Science 334, 369–373 (2011).

16. Shehzad, Z. et al. An Multivariate Distance-Based Analytic Framework for Connectome-Wide Association Studies. NeuroImage 93, 74–94 (2014).

17. Kuang, D., Ding, C. & Park, H. Symmetric Nonnegative Matrix Factorization for Graph Clustering. in Proceedings of the 2012 SIAM International Conference on Data Mining 106–117 (Society for Industrial and Applied Mathematics, 2012). doi:10.1137/1.9781611972825.10

18. Ray, K. L. et al. ICA model order selection of task co-activation networks. Front. Neurosci. 7, (2013).

19. Bolt, T., Nomi, J. S., Yeo, B. T. T. & Uddin, L. Q. Data-Driven Extraction of a Nested Model of Human Brain Function. J. Neurosci. 37, 7263–7277 (2017).

20. Stroop, J. Studies of interference in serial verbal reactions. J. Exp. Psychol. 18, 643–662 (1935).

21. Friedman, L. et al. Brain Activation During Silent Word Generation Evaluated with Functional MRI. Brain Lang. 64, 231–256 (1998).

22. Eklund, A., Nichols, T. E. & Knutsson, H. Cluster failure: Why fMRI inferences for spatial extent have inflated false-positive rates. Proc. Natl. Acad. Sci. 113, 7900–7905 (2016).

23. Gorgolewski, K. J. et al. NeuroVault.org: a web-based repository for collecting and sharing unthresholded statistical maps of the human brain. Front. Neuroinformatics 9, 8 (2015).

24. Fox, M. D. et al. The human brain is intrinsically organized into dynamic, anticorrelated functional networks. Proc. Natl. Acad. Sci. U. S. A. 102, 9673–9678 (2005).

25. Raichle, M. E. et al. A default mode of brain function. Proc. Natl. Acad. Sci. 98, 676–682 (2001).

26. Uddin, L. Q. Salience processing and insular cortical function and dysfunction. Nat. Rev. Neurosci. 16, 55–61 (2015).

27. Lancaster, J. L. et al. Bias between MNI and Talairach coordinates analyzed using the ICBM-152 brain template. Hum. Brain Mapp. 28, 1194–1205 (2007).

28. Kim, J. & Park, H. Fast Nonnegative Matrix Factorization: An Active-Set-Like Method and Comparisons. SIAM J. Sci. Comput. 33, 3261–3281 (2011).

29. Boutsidis, C. & Gallopoulos, E. SVD based initialization: A head start for nonnegative matrix factorization. Pattern Recognit. 41, 1350–1362 (2008).

30. Sotiras, A., Resnick, S. M. & Davatzikos, C. Finding Imaging Patterns of Structural Covariance via Non-Negative Matrix Factorization. NeuroImage 108, 1–16 (2015).

31. Dhillon, I. S. & Modha, D. S. Concept Decompositions for Large Sparse Text Data Using Clustering. Mach. Learn. 42, 143–175 (2001).

32. Shirkhorshidi, A. S., Aghabozorgi, S. & Wah, T. Y. A Comparison Study on Similarity and Dissimilarity Measures in Clustering Continuous Data. PLOS ONE 10, e0144059 (2015).

33. Wu, S. et al. Stability-driven nonnegative matrix factorization to interpret spatial gene expression and build local gene networks. Proc. Natl. Acad. Sci. 113, 4290–4295 (2016).

34. Garcia, D. Robust smoothing of gridded data in one and higher dimensions with missing values. Comput. Stat. Data Anal. 54, 1167–1178 (2010).

